# Sustained modulation of emotion-related fibers after 8 weeks of mindfulness-based stress reduction training

**DOI:** 10.1101/855551

**Authors:** Chang-Le Chen, Yao-Chia Shih, Yung-Chin Hsu, Tzung-Kuen Wen, Shih-Chin Fang, Da-Lun Tang, Si-Chen Lee, Wen-Yih Isaac Tseng

## Abstract

An 8-week mindfulness-based stress reduction (MBSR) program enables novices to learn and practice mindfulness meditation. A longitudinal study design has been used in prior research to investigate the effect of short-term mindfulness meditation training on brain structures. Many studies have demonstrated microstructural changes in the white matter by comparing baseline measurements with measurements obtained immediately after short-term meditation training. However, these studies did not clarify the evolution of the modulated microstructures several months after mindfulness meditation practice is discontinued. Therefore, in this study, we recruited 13 novice practitioners and administered an 8-week MBSR training program. We extended the span of the longitudinal study by adding a third measurement taken approximately 6 months after the second time point. Diffusion indices derived from diffusion spectrum imaging were used to quantify the temporal changes in modulation across three time points. The analysis identified four tract bundles that were significantly modulated after the 8-week MBSR training program, namely the callosal fibers connecting the bilateral amygdalae and bilateral hippocampi, right thalamic radiation of the auditory nerve, and right uncinate fasciculus. At the third time point, at which the participants had discontinued practice for approximately 6 months, the diffusion indices of the four tract bundles still presented a significant difference compared with the baseline. Our results indicate that the modulation of microstructural properties of the white matter tract induced by the 8-week MBSR program was sustained after completion of the program and support that neuroplasticity in brain connection persists after the discontinuation of meditation training.

## Introduction

Mindfulness meditation originates from ancient Buddhist meditation traditions (Kabat-Zinn 2003) and is usually described as nonjudgmental attention to the present-moment experience (Chiesa and Malinowski 2011). Mindfulness meditation practice encompasses focusing on the experience of thoughts, sensations, and emotions and simply observing them as they occur (Holzel et al. 2011a). Mindfulness-based stress reduction (MBSR) is a modern standardized mindfulness meditation training program that is conducted over a period of 8 weeks, in which participants are instructed to connect their physical sensations, perceptions, and emotions through body awareness, yoga, and mindfulness meditation (Kabat-Zinn and Hanh 2009). This 8-week program is considered useful for reducing stress (Chiesa and Serretti 2009), anxiety (Goldin et al. 2009; Goldin and Gross 2010), and depression (Sephton et al. 2007) and improving self-awareness (Shapiro et al. 2007), attention (Anderson et al. 2007; Jha et al. 2007), and interoceptive sensitivity (Melloni et al. 2013). Because of the psychological benefits manifested through MBSR, considerable attention has been paid to the investigation of the psychological and underlying neurological mechanisms of MBSR.

Potential neuroanatomical changes caused by mindfulness meditation have been investigated in prior research. Changes in the cortical or subcortical brain structures have been proven to underpin the neuroplasticity induced by mindfulness meditation (Holzel et al. 2011b; Lazar et al. 2005; Santarnecchi et al. 2014). A neuroimaging study demonstrated increased regional gray matter density in practitioners who performed mindfulness meditation compared with naïve controls (Holzel et al. 2011b). In addition to the effects of MBSR on gray matter, several studies have evidenced improved anatomical connectivity of the white matter in the large-scale dynamic network of high-level cognition induced by mindfulness meditation (Kang et al. 2013; Luders et al. 2011, 2012). Under normal circumstances, the white matter responds to experiences that affect brain function, thereby altering performance and information processing (Mackey et al. 2012). Mackey et al. (2012) described how learning and experiences shape the microstructure of white matter. Using diffusion tensor imaging (DTI), Luders et al. (2011) found pronounced structural connectivity throughout the entire brain in long-term meditators compared with controls. In another study by Luders et al. (2012), experienced practitioners exhibited thickening of the corpus callosum with increased values of fractional anisotropy (FA), which is a diffusion index derived from DTI to indicate microstructural integrity (Basser and Pierpaoli, 2011; Beaulieu 2002). These findings imply that increased interhemispheric integration is associated with characteristic mental states and meditation skills. However, the aforementioned studies and most other meditation-related neuroimaging reports have adopted a cross-sectional study design to compare meditators and controls (Fox et al. 2014). Because these studies were cross-sectional, they could not clarify whether the observed structural difference was actually caused by meditation or whether the difference was preexisting and thus made a specific group of people more likely than others to engage in intensive meditation practices (Tang et al. 2015).

Recently, more studies adopted a longitudinal design to investigate the neuroanatomical changes induced by short-term mindfulness meditation. This type of approach can demonstrate a direct causal relationship of this nature and is often conducted in meditation-naïve novices who receive short-term meditation training. In this design, usually known as the pre–post intervention style, brain structures are examined before and after training and the differences are compared between novice practitioners and control participants. Previous longitudinal experiments have determined structural differences in novice practitioners after short-term meditation training. Tang et al. (2010) reported that short-term integrative body– mind training (IBMT) increased the FA values of the white matter tracts connecting the anterior cingulate cortex. Moreover, they proved that the changes in white matter connectivity can be observed after a short-term intensive mental training program. Previous studies that adopted the pre–post experimental design to investigate the microstructural changes of the white matter reported a rapid induction of alterations in the brain structure. However, it raises the question of whether such a change disappears with equal speed in the absence of continual practice (Fox et al. 2014). For example, gray matter structure changes caused by motor skill learning have been demonstrated to diminish without continual practice (Draganski et al. 2004). This type of follow-up study, despite being difficult to execute, is an ideal paradigm for clarifying the speed at which neuroanatomical changes wane after the cessation of intensive meditation practices. A potential implication is that neuroplasticity is induced by short-term mental training, providing a rationale for clinical mindfulness-related cognitive therapy.

The aim of the present study was to clarify whether the neuroanatomical changes induced by short-term mindfulness meditation was transient or sustained. Specifically, we investigated change in the microstructural properties of the white matter tracts by conducting two experiments. The first experiment was an 8-week longitudinal study in which we compared the microstructural change induced by short-term mindfulness meditation training in the whole-brain white matter tracts between meditation novices and controls. The MBSR program was used to train the novices. In the second experiment, we extended the 8-week longitudinal design by monitoring the training-induced microstructural changes approximately 6 months after the program ended. All novices were required not to practice MBSR during this period. We ascertained whether the changes persisted or diminished after the cessation of practice, hypothesizing that the changes induced by short-term mindfulness meditation training would be sustained up to the third time point.

## Materials and methods

### Experimental design

In the present study, we adopted a three-time-point longitudinal design (Fig. 1) to determine whether the microstructural changes induced by the 8-week MBSR program were transient or sustained if the practitioners ceased to practice meditation for a prolonged period. Experiment 1 entailed an 8-week longitudinal study (Fig. 1, left) comprising the MBSR and control groups. This experiment explored the microstructural alterations induced by short-term meditation training. The MBSR group underwent an initial magnetic resonance imaging (MRI) scan 1 month before the MBSR program (time point 1, TP1) and the second scan within 1 month after the end of the program (time point 2, TP2). The participants in the control group underwent their first and second scans 8 weeks apart with no mental training during this period. Experiment 2 involved a 6-month follow-up study (Fig. 1, right) performed on the MBSR group only. This experiment determined the transience or sustainability of the microstructures that were modulated by the MBSR program. The third scan was performed approximately 6 months (mean/SD: 6.1/2.1 months) after completion of the program (time point 3, TP3). All participants were instructed to discontinue any MBSR practice during this period. A self-report was obtained from the participants to confirm that they did not practice MBSR.

**Fig. 1.**
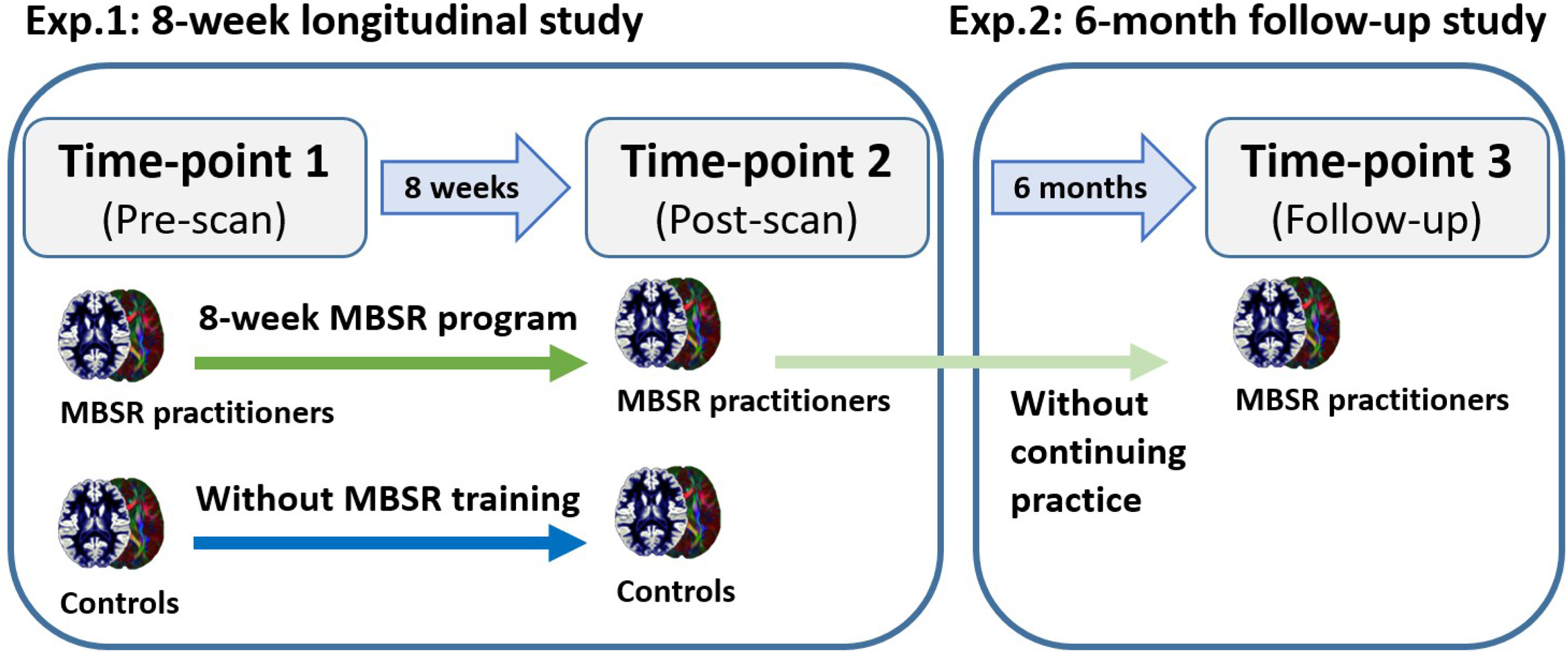
Experimental design of the three-time-point longitudinal study. Experiment 1 (Exp. 1) entailed an 8-week longitudinal study analyzing the MBSR and control groups. All participants underwent two MRI scans; the MBSR group underwent one scan before the MBSR program and one after and the control group underwent two scans 8 weeks apart without having received any mental training. Experiment 2 (Exp. 2) entailed a follow-up study approximately 6 months after the completion of the MBSR program. The practitioners were required to refrain from any MBSR practice during this period.

### Participants

Initially, 30 participants were recruited from respondents to an advertisement for an 8-week MBSR program promoted by the Taiwan Mindfulness Development Association. All participants were Taiwanese, right-handed, and declared no previous experience of mindfulness and meditation practices. Participants with a previous history of neurological or psychiatric disorders or contraindications for MRI scanning were excluded. Only 15 participants completed the entire 8-week course and two time point scans. For the follow-up study, 13 out of the 15 participants completed all the MRI scans over three time points. Finally, 13 participants (*n* = 13; five men; mean age: 41.3 ± 7.6 years) with three time point examinations were assigned to the MBSR group, and an additional 13 participants (*n* = 13; 3 men; mean age: 42.0 ± 15.4 years) were recruited as the control group. Notably, the control group was scanned at only two time points during the same time period of the MBSR program. The two groups did not significantly differ in mean age (t_(24)_ = 0.14, *p* = 0.89) and sex distribution (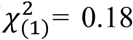, *p* = 0.67). In the longitudinal study, the stability of measurement system is of importance for repeated measures (details seen in section 2.8). To assess the test– retest reliability of the MRI system, another group of 26 healthy adults (four men; mean age: 52.8 ± 15.6 years) was recruited. The participants in this group underwent two MRI scans approximately 3 months apart, without any intervention applied during this period. All examinations were approved by the Institutional Review Board of the National Taiwan University Hospital, and informed consent was obtained from each participant.

### Training methods

The MBSR program (Kabat-Zinn 1990) has been promoted worldwide over the past two decades (Grossman et al. 2004). We adopted a paradigm that included an 8-week group course with 3-h sessions on Saturdays and individual practice for 1–2 h every day from Sunday through Friday. Mindfulness training was imparted by a certified instructor (T.K.W.) to develop the psychological capacity for mindfulness (i.e., awareness of present-moment experiences with a nonjudgmental stance; Germer 2004). The course content included a body scan, mindful yoga, and sitting meditation. During the body scan process, attention was guided sequentially throughout the entire body to observe the sensations in each region without judgment. Mindful yoga entailed gradual movements and gentle outstretching exercises that acted in concert with breathing. Sitting meditation began with the awareness of breathing and evolved to integrate different modalities of awareness such as sounds and emotions. To ensure that the MBSR participants practiced regularly during the 8-week MBSR program, they were requested to report home practice compliance and record the amount of time spent on mindfulness exercises each day. Participants enrolled in the MBSR group were considered eligible for the study if they completed the MBSR formal course and daily assignments.

### Psychological assessments

To confirm that the MBSR participants had been trained and instructed effectively, their mindfulness-related psychological properties were evaluated using the Chinese version of the five-facet mindfulness questionnaire (Ch-FFMQ) (Deng et al. 2011) at each time point. Originally developed by Baer et al. (2006), this questionnaire is used to assess an individual’s trait-like general tendency toward mindfulness. The FFMQ is a self-report instrument comprising 39 items and a 5-point Likert-type scale for assessing five identified facets of mindfulness: nonjudgment of inner experience, nonreactivity to inner experience, acting with awareness, describing, and observing. The Cronbach’s α values and Guttmann split-half coefficients for the five facets of mindfulness scores in Ch-FFMQ were 0.698 and 0.679 on average, respectively, indicating adequate reliability and internal consistency (Deng et al. 2011). A paired *t* test was used for hypothesis testing to compare the pre–post differences.

### Image acquisitions

The microstructural properties of the white matter were measured using various diffusion indices derived from diffusion spectrum imaging (DSI). DSI is a diffusion MRI technique developed for mapping spatial profiles of the diffusion average propagator and has been demonstrated to resolve crossing fibers (Jonasson et al. 2005; Wedeen et al. 2005). All images were acquired on a 3T MRI system (Tim Trio; Siemens, Erlangen, Germany) with a 32-channel phased-array head coil. All participants lay on the MRI table with their heads packed with expandable foam cushions to restrict movement. To obtain an anatomical reference, high-resolution T1-weighted imaging was performed using a 3D magnetization-prepared rapid gradient echo sequence: repetition time (TR) / echo time (TE) = 2000/3 ms, flip angle = 9°, field of view (FOV) = 256 × 192 × 208 mm^3^, and acquisition matrix = 256 × 192 × 208, resulting in an isotropic spatial resolution of 1 mm^3^. DSI was performed using a pulsed-gradient spin-echo diffusion echo planar imaging sequence with a twice-refocused balanced echo (Reese et al. 2003). The imaging parameters were as follows: b_max_ = 4000 s/mm^2^, TR/TE = 9600/130 ms, slice thickness = 2.5 mm, acquisition matrix = 80 × 80, FOV = 200 × 200 mm^2^, and in-plane spatial resolution = 2.5 × 2.5 mm^2^. The diffusion-encoding acquisition scheme used in the present study followed the framework of DSI (Wedeen et al. 2005), to which 102 diffusion-encoding gradients were applied corresponding to the grid points in the half-sphere of the 3D q-space within a radius of three units (Kuo et al. 2008). Because the data in the q-space are real and symmetrical around the origin, the acquired half-sphere data were projected to fill the other half of the sphere. Each scan, including T1-weighted imaging and DSI, was completed within approximately 20 minutes.

### Image analyses

#### Image quality assurance

DSI data sets were subjected to quality assurance procedures, including examinations of signal-to-noise ratio (SNR) and in-scanner head motion. SNR was evaluated by calculating the mean signal of an object divided by the standard deviation of the background noise (Dietrich et al. 2007). In practice, the signal was determined using a central square of an image with a size of 20 × 20 pixels, and the noise was averaged from four corner regions with a size of 10 × 10 pixels for each region. DSI data sets with an averaged SNR of <20 were discarded. Because of the relatively long scan time required for DSI acquisition, in-scanner head motion is inevitable and may cause signal dropout in diffusion-weighted images, particularly in those with high b values. All acquired DSI data sets (5712 images per participant) were examined by comparing the signal in the central square of each image with the predicted signal attenuation. Signal deviation from the predicted distribution was considered as signal loss. Data with more than 60 images of signal dropout per participant (1% of the total diffusion-weighted images) were discarded. All data used in this study passed the quality assurance criteria.

#### Diffusion index calculation

The diffusion indices derived from DSI were computed according to the framework of mean apparent propagator (MAP)-MRI (Ozarslan et al. 2013). MAP-MRI provides a quantitative, efficient, and robust mathematical framework for modeling a signal in a three-dimensional q-space, which is the Fourier conjugate of the diffusion propagator. The signal in the q-space was fitted with a series expansion of Hermite basis functions, which accurately describe diffusion in various microstructural geometries (Avram et al. 2016). The lowest order term in the expansion contains a diffusion tensor that characterizes the Gaussian displacement distribution. The higher-order terms in the expansion are the orthogonal corrections to the Gaussian approximation and enable the reconstruction of the average propagator.

Diagonalization of the diffusion tensor provides eigenvalues and the corresponding eigenvectors. The values of axial diffusivity (AD), radial diffusivity (RD), and mean diffusivity (MD) at each voxel were determined using the eigenvalues of the diffusion tensor model. The AD, RD, and MD were quantified as the first eigenvalue, mean of the second and third eigenvalues, and mean of the three eigenvalues, respectively. Orientation distribution function (ODF) was defined as the second radial moment of the average propagator and was analytically computed from the series coefficients of Hermite functions. The generalized FA (GFA) value was quantified as the standard deviation of ODF divided by the root mean square of ODF (Fritzsche et al. 2010; Gorczewski et al. 2009). Similar to the FA, GFA provides the anisotropic diffusion information of water molecules, which may reflect the microstructural architecture of the white matter. The diffusion indices (i.e., GFA, AD, MD, and RD) represent multiple aspects of the microstructural properties of the white matter, such as degree of myelination, fiber calibers, and fiber density (Alexander et al. 2011).

#### Whole-brain tract-specific analysis

Whole-brain tract-based automatic analysis (TBAA) was conducted to investigate tract-specific differences in the microstructural properties between meditation novices and controls (Chen et al. 2015). TBAA enables automatic tract-specific analysis of the 76 major fiber tract bundles over the whole brain. The TBAA was used to generate a DSI template called NTU-DSI-122 (Hsu et al. 2015) and a table of tract coordinates in the template to sample the diffusion indices (i.e., GFA, AD, RD, and MD) of each tract from each individual’s diffusion data set through the known transformation between the DSI template and individual DSI data set.

NTU-DSI-122 is a high-quality DSI template averaged over 122 coregistered DSI data sets of healthy adults and built in the standard ICBM-152 space. The 76 tract bundles were segmented in the template by applying deterministic streamline-based tractography with multiple regions of interest defined in the automated anatomical labeling atlas (Tzourio-Mazoyer et al. 2002). The coordinates of the streamlines were aligned with the proceeding direction of each tract bundle, normalized into 100 steps, and recorded as the sampling coordinates in the template space (Chen et al. 2015).

The overall procedure of the TBAA method is presented in Fig. 2 and described as follows. A total of 65 DSI data sets (13 MBSR participants at three time points and 13 control participants at two time points) were registered using a two-step strategy to construct a study-specific template (SST), which was then registered to the NTU-DSI-122 template through the same strategy. The two-step registration method was modified for diffusion data sets (Hsu et al. 2012). The strategy harnessed anatomical information obtained from T1-weighted images and microstructural information obtained from DSI data sets. The first step was to register the individual tissue probability map (TPM) from the T1-weighted images into mean TPM and then to transform the individual DSI data sets into the mean TPM space to build an intermediate DSI template. The second step was to normalize the registered DSI data sets to the intermediate template by performing DSI-specific large deformation diffeomorphic metric mapping (Hsu et al. 2012). Once the registration procedure was completed, the sampling coordinates of the predefined 76 major white matter tracts were transformed from the standard template space to individual DSI data sets through the transformation processes between the NTU-DSI-122 and SST and between the SST and individual DSI data sets. Diffusion indices were sampled in the individual DSI space by using the coordinates transformed from the standard space. The output of TBAA was a 2D matrix of the diffusion index for each participant, called the connectogram. Each row of the connectogram indicated the values of the diffusion index sampled on 100 steps along each tract bundle. A 3D connectogram was formed when multiple 2D connectograms of the study participants were stacked together. Before statistical analyses were performed, each row of the connectogram was spatially smoothed by convolving a Gaussian kernel with a full-width half-maximum of 20 steps to reduce local variance. Due to zero-padding in the convolution procedure, we discarded five steps at each end of the row, thus reducing the number of steps from 100 to 90. Finally, we obtained four 3D connectograms of GFA, AD, MD, and RD from the MBSR and control groups.

**Fig. 2.**
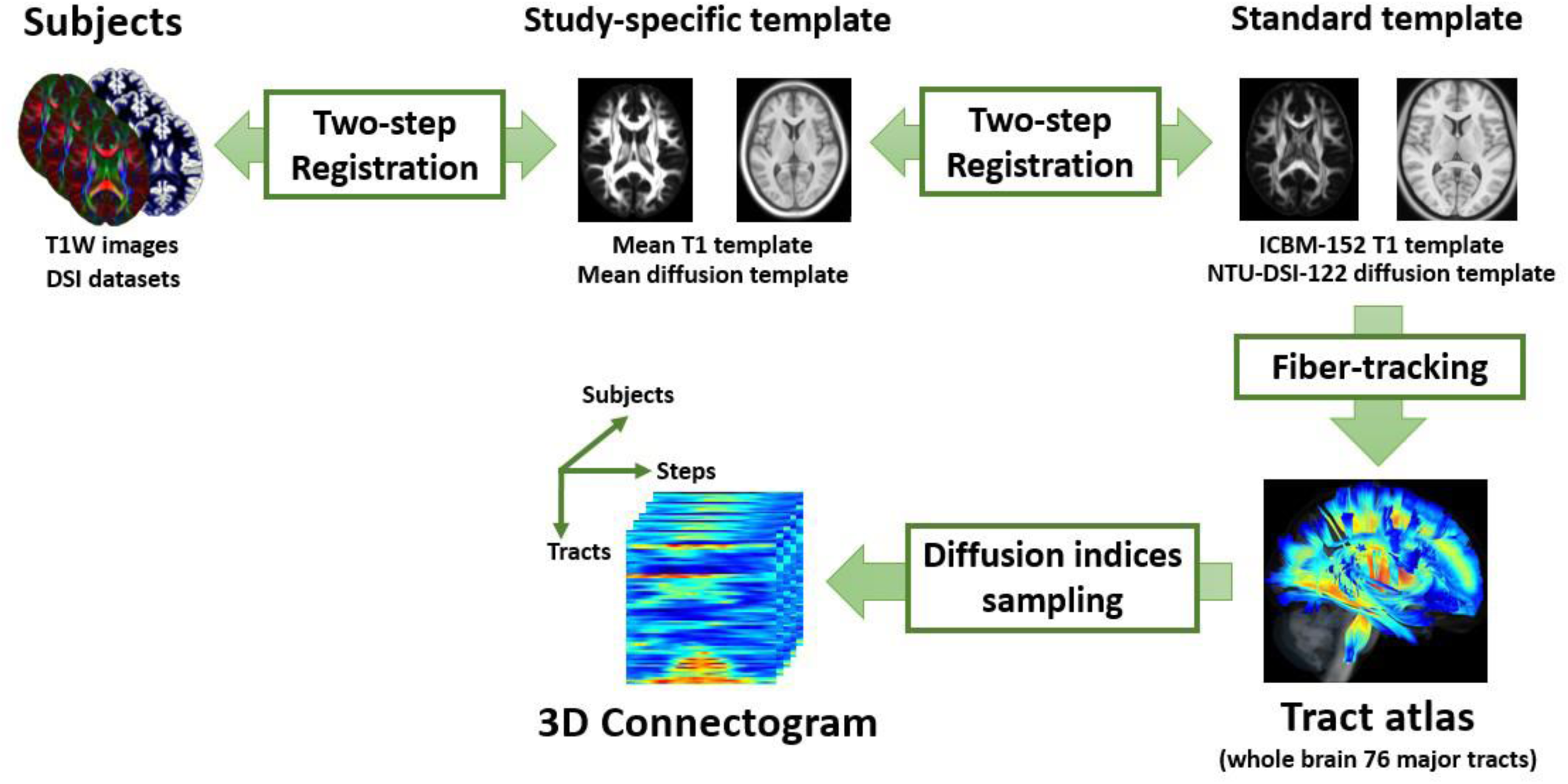
Flowchart of the TBAA procedures.

### Statistical analyses of imaging data

In the 8-week longitudinal study (Experiment 1), a cohort analysis was first conducted to exclude potential prior differences between the groups. We compared the MBSR and control groups at the first time point by using a two-sample *t* test at each step of the connectogram. Multiple-comparison correction was performed using a cluster threshold: cluster size connection > 15 steps and individual step threshold, *p* < 0.01. In the longitudinal analysis, four repeated measures analysis of covariance (RM-ANCOVA) models were performed for four diffusion indices (i.e., GFA, AD, RD, and MD) by using R-Studio (version 1.0) and the Statistical and Machine Learning Toolbox for MATLAB (version 9.1). RM-ANCOVA was used to test the differences in each diffusion index at each step of the connectogram, with “Group” (MBSR and wait-list controls) as the between-subject factor and “Pre–Post” (the first and second MRI examination) as the within-subject factor. Age and sex were included as covariates. Multiple-comparison correction was performed using the same threshold criteria as those for the cohort analysis to eliminate false discovery. The significant primary effect was followed by post hoc comparisons to examine the within-factor “Time” differences for each group, where *p* < 0.05 was considered statistically significant. In addition, we calculated the effect size—Cohen’s d_rm_, an equivalent of Cohen’s d for repeated measures (Lakens 2013; Morris and DeShon 2002)—of the simple effect “Time” in each group for quantifying substantive significance (Sullivan and Feinn 2012).

In the 6-month follow-up study (Experiment 2), the statistically altered segments obtained in the 8-week longitudinal study were selected to examine the changes across three time points. Repeated measures analysis of variance (RM-ANOVA) was performed as the follow-up hypothesis test on those segments, and TP1, TP2, and TP3 were used as the within-subject factors. Bonferroni correction was used to address the multiple-comparison problem. The primary effect was followed by post hoc comparisons to examine the differences between TP2 and TP1 and between TP3 and TP1. The differences in diffusion indices were quantified as percent changes calculated using the following formula:

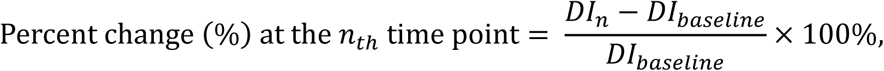

where DI_n_ and DI_baseline_ indicates the diffusion indices at the *n_th_* time point and baseline (TP1), respectively.

### System repeatability assessments

A longitudinal study requires repeated measures and is highly dependent on the stability of the system. The microstructural changes induced by short-term training could be so subtle as to be overwhelmed by the bias of the measuring system. Therefore, the intraclass correlation coefficient (ICC) was applied to assess the test-retest reliability, also called repeatability, of the MRI system (Weir 2005). A total of 26 healthy adults were recruited to undergo two MRI scans approximately 3 months apart. The MRI data, including T1-weighted images and DSI data sets, were analyzed using TBAA to generate connectograms with the aforementioned procedures. Repeatability was evaluated stepwise on the two GFA connectograms of the same participant by using the second formula of ICC (Shrout and Fleiss 1979). Overall repeatability was determined by averaging the repeatability values over all steps of the connectogram.

## Results

### System repeatability assessments

The repeatability was analyzed on all 26 participants, and the mean ICC was 0.931 ± 0.0418, indicating the high test-retest reliability and ensuring that the measurement system was stable and reliable. (Fig. 3).

**Fig. 3.**
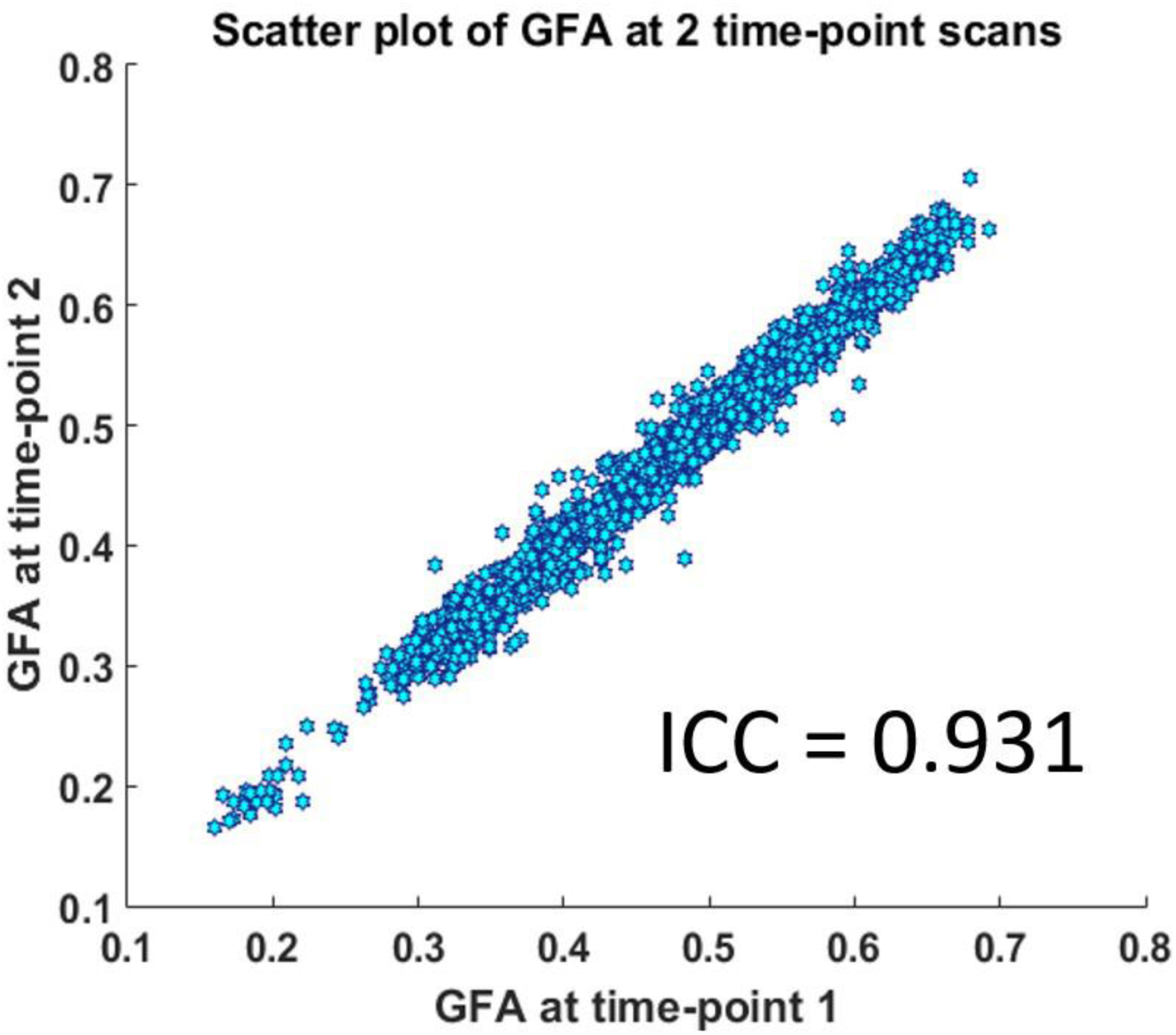
Scatter plot of the GFA values at two time points. X and Y axes indicate the GFA values measured at time points 1 and 2, respectively.

### Psychological assessments

The MBSR group was evaluated using the Ch-FFMQ to assess mindfulness-related psychological improvements following the 8-week MBSR program. The paired *t* test revealed significant increases in four of the five mindfulness subscales (observe: *t*_(12)_ = 3.02, *p* = 0.011; describe: *t*_(12)_ = 2.45, *p* = 0.030; act with awareness: *t*_(12)_ = 3.01, *p* = 0.011; nonjudgment: *t*_(12)_ = 1.45, *p* = 0.173; and nonreactivity: *t*_(12)_ = 2.40, *p* = 0.034). The total score on the Ch-FFMQ also significantly increased after the MBSR program (*t*_(12)_ = 4.17, *p* = 0.001). In sum, the MBSR participants exhibited significantly improved mindfulness scores in all facets except the nonjudgment scale (Fig. 4).

**Fig. 4.**
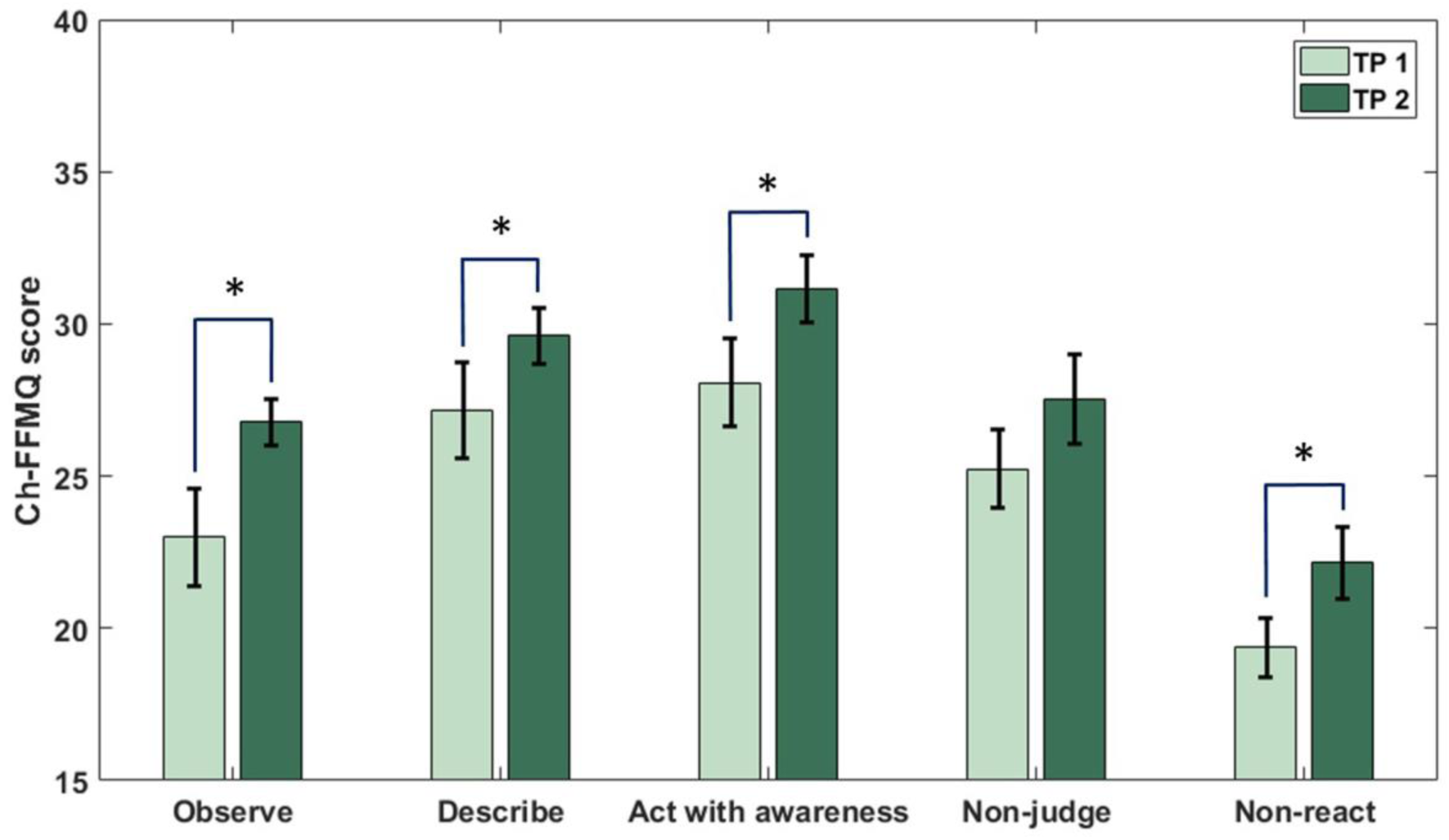
Significant improvement in Ch-FFMQ scores after the 8-week MBSR program, except in the “nonjudgment” facet. TP1 and TP2 denote time point 1 (baseline) and time point 2 (after the MBSR program), respectively. The asterisk indicates statistical significance, and the error bar indicates one standard error.

### Eight-week longitudinal analysis (Experiment 1)

Cohort analysis was performed at the baseline to identify any preexisting differences between the MBSR and control groups. No significant differences were found in GFA, AD, MD, or RD between the groups. In the 8-week longitudinal analysis, RM-ANCOVA modeling was performed on GFA, AD, RD, and MD values separately, with age and sex as covariates. In the GFA connectogram, the interaction between “Group” and “Time” was significantly different in three white matter tract segments (i.e., the right thalamic radiation [TR] of the auditory nerve [peak: *F*_(1, 22)_ = 20.1; *p* = 0.0002, segment: 35 steps], the callosal fibers [CF] connecting the bilateral hippocampi [peak: *F*_(1, 22)_ = 16.4; *p* = 0.0005, segment: 16 steps], and the CF connecting the bilateral amygdalae [peak: *F*_(1, 22)_ = 12.8; *p* = 0.0017, segment: 17 steps]) (Table 1). The following post hoc comparisons of the primary effect indicated that the segments significantly increased (*p* < 0.05) in the MBSR participants from TP1 to TP2, whereas no significant difference (*p* > 0.05) was observed in the control group. In the subsequent effect size calculation, Cohen’s d_rm_ was 1.11, 0.69, and 0.59 for the right TR of the auditory nerve, CF of the hippocampi, and CF of the amygdalae, respectively. These segments exhibited a significant increase in GFA values after the 8-week MBSR program (Fig. 5). In both the AD and MD connectograms, the interaction between “Group” and “Time” was significantly different in the segments of one white matter tract (i.e., the right uncinate fasciculus [UF] [peak: *F*_(1, 22)_ = 15.7; *p* = 0.0007, segment: 21 steps in AD, peak: *F*_(1, 22)_ = 12.7; *p* = 0.0017, segment: 16 steps in MD]). The post hoc comparisons of the primary effect revealed that the segments significantly decreased in the MBSR participants from TP1 to TP2, whereas no significant difference was observed in the control group (Table 1). In the MBSR group, the Cohen’s d_rm_ in the segments of the right UF was 0.93 and 0.56 for AD and MD, respectively. These segments exhibited a significant decrease in AD and MD values after the 8-week MBSR program (Fig. 6). No significant difference was observed in the statistical interaction in RD.

**Fig. 5.**
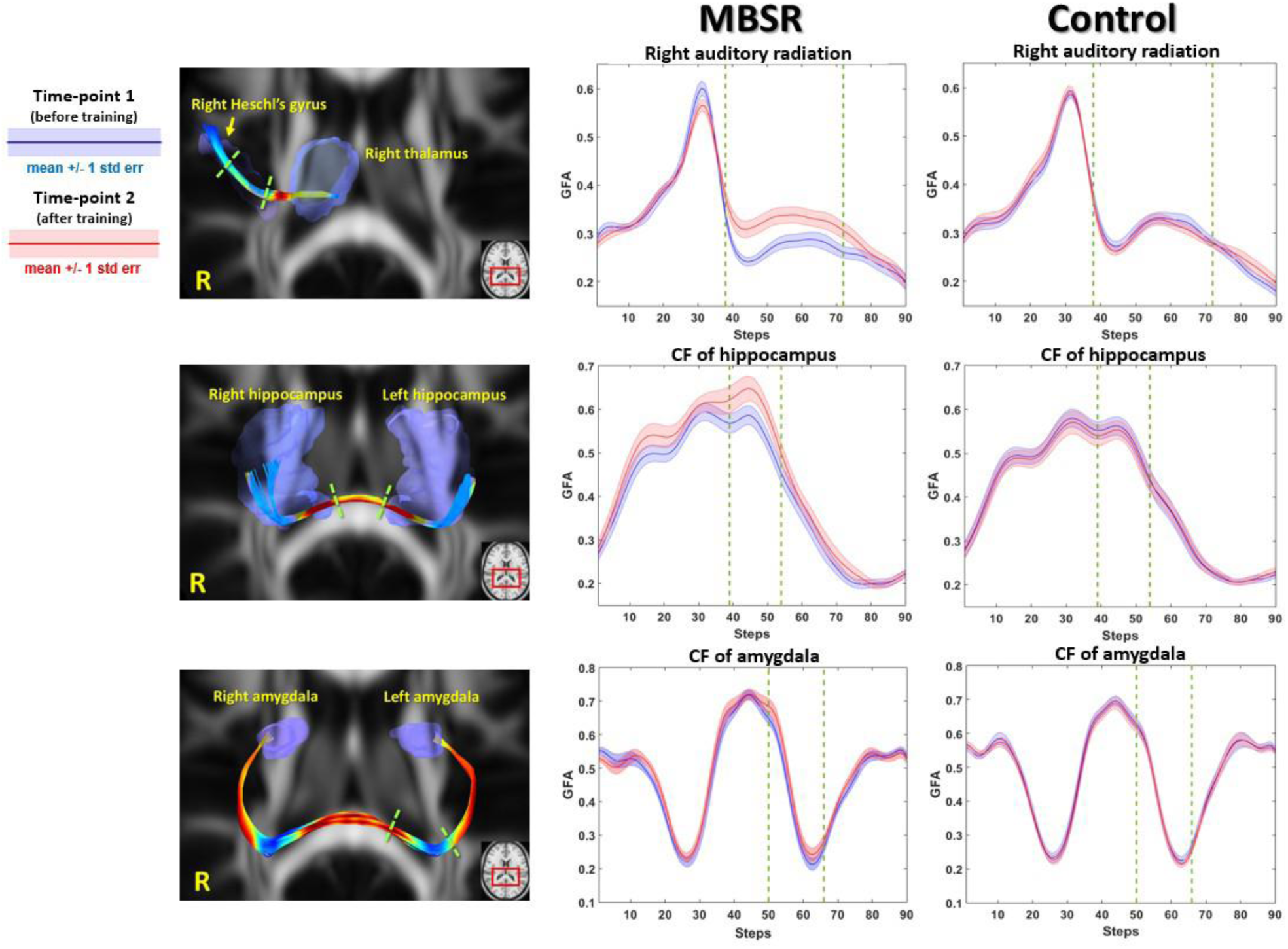
White matter segments exhibiting a significant increase in GFA following short-term MBSR training in the MBSR group. These segments include the right TR of the auditory nerve (top), CF of the hippocampus (middle), and CF of the amygdala (bottom). The left column displays the fiber pathways of the tracts and the significantly different segments, which are bounded by two dashed green lines. The middle and right columns present the tract profiles of the MBSR and control groups, respectively. The segments bounded by the two dashed green lines exhibit a significant difference. Blue lines and shaded error bars represent the mean response and one standard error at the baseline (time point 1), respectively. Red lines and shaded error bars represent the mean response and one standard error after the MBSR program (time point 2), respectively.

**Fig. 6.**
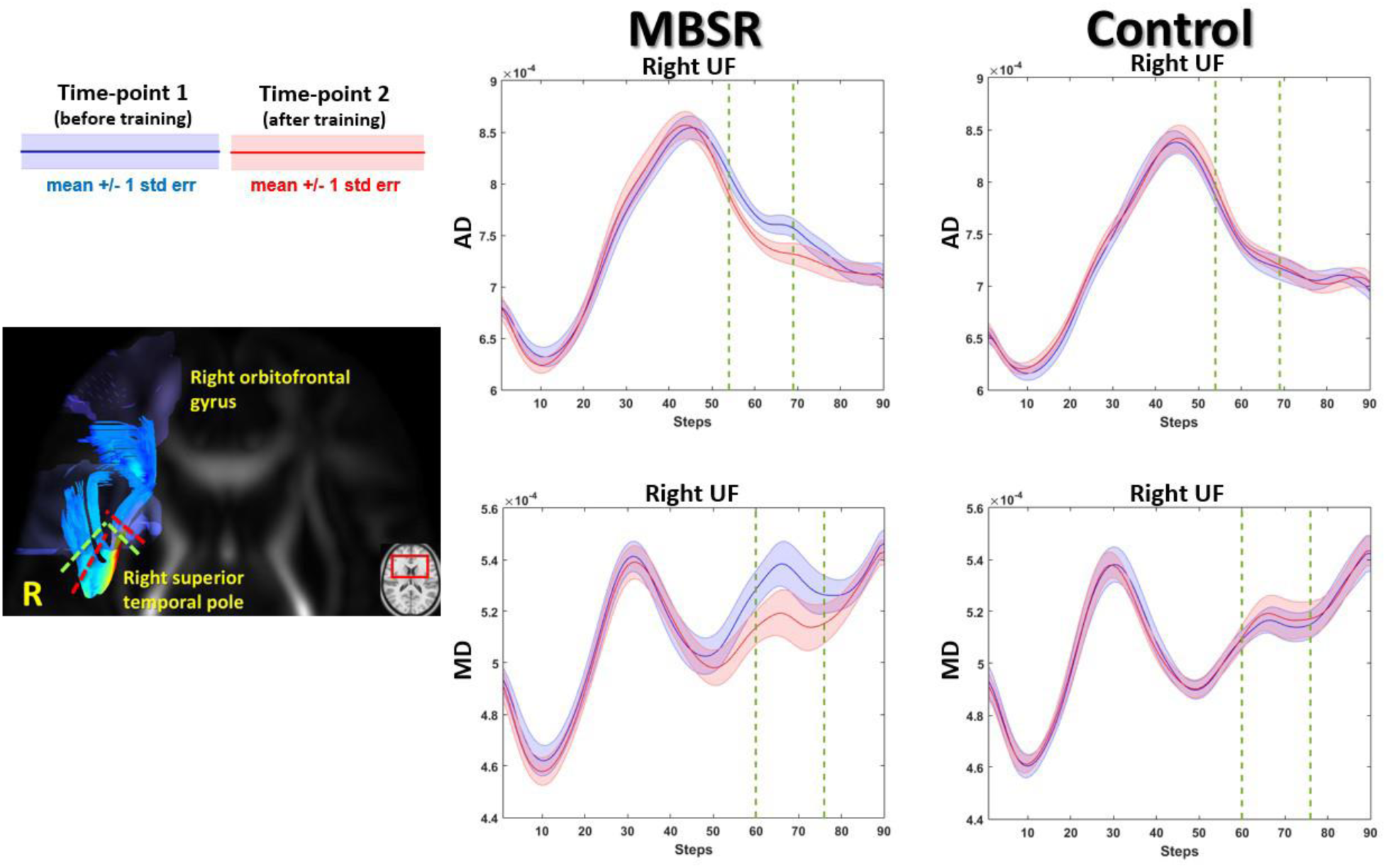
Significant decrease in AD and MD in the right UF following short-term MBSR training in the MBSR group. The tract profiles in the MBSR and control groups are shown in the middle and right columns, respectively. The segments bounded by the two dashed green lines exhibit a significant difference. In the fiber pathways of the right UF (left panel), the segments bounded by the red and green dash lines exhibit a significant difference in AD and MD, respectively. Lines and shaded error bars are presented in the same manner as in Fig. 5.

**Table 1.** Results of repeated measures ANCOVA in the 8-week longitudinal study. d_rm_ indicates the effect size Cohen’s d_rm_ for repeated measures in the simple effect “Time”

| Diffusion indices | Tract | MBSR group |  | Control group |  | Main effect |  | Simple effect (Time) |  |  |  | Cluster size |
| --- | --- | --- | --- | --- | --- | --- | --- | --- | --- | --- | --- | --- |
|  |  | Time-point 1 | Time-point 2 | Time-point 1 | Time-point 2 | Group × Time |  | MBSR |  | Control |  |  |
|  |  | Mean (SD) | Mean (SD) | Mean (SD) | Mean (SD) | *F* (peak) | *p* | *p* | *d*<sub>rm</sub> | *p* | *d*<sub>rm</sub> |  |
| GFA | R TR of auditory nerve | 0.273 (0.040) | 0.328 (0.054) | 0.310 (0.035) | 0.302 (0.028) | 20.1 | < 0.01 | < 0.05 | 1.11 | > 0.05 | 0.26 | 35 |
|  | CF of hippocampus | 0.547 (0.073) | 0.606 (0.090) | 0.530 (0.074) | 0.521 (0.077) | 16.4 | < 0.01 | < 0.05 | 0.69 | > 0.05 | 0.12 | 16 |
|  | CF of amygdala | 0.381 (0.052) | 0.417 (0.063) | 0.381 (0.049) | 0.375 (0.052) | 12.8 | < 0.01 | < 0.05 | 0.59 | > 0.05 | 0.10 | 17 |
| AD | R UF<sup>+</sup> | 7.729 (0.257) | 7.498 (0.237) | 7.406 (0.307) | 7.459 (0.314) | 15.7 | < 0.01 | < 0.05 | 0.93 | > 0.05 | 0.17 | 21 |
| MD | R UF<sup>+</sup> | 5.327 (0.291) | 5.162 (0.295) | 5.142 (0.141) | 5.166 (0.225) | 12.7 | < 0.01 | < 0.05 | 0.56 | > 0.05 | 0.11 | 16 |

<sup>+</sup>: ( x 10<sup>-4</sup> ) mm<sup>2</sup>/sec unit

### Six-month follow-up study (Experiment 2)

The third MRI scan was performed on the MBSR group approximately 6 months after completion of the MBSR program. The follow-up analysis was performed on the segments that showed significant modulation during the 8-week longitudinal study. In the repeated measures ANOVA, Mauchly’s test indicated that the assumption of sphericity was not violated, except in the CF of the hippocampi (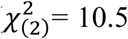, *p* = 0.005) and CF of the amygdalae (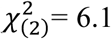, *p* = 0.048); therefore, the Greenhouse–Geisser correction was used to address the violation of sphericity (Table 2). The GFA values of the segments in the right TR of the auditory nerve (*F*_(2, 24)_ = 11.1; *p* = 0.0004), CF of the hippocampi (*F*_(1.2, 14.9)_ = 13.0; *p* = 0.0017), and CF of the amygdalae (*F*_(1.4, 16.9)_ = 13.1; *p* = 0.0010) were significantly different over three time points. In the post hoc analysis, all the GFA values of the segments significantly increased from TP1 to TP2 (the right TR of the auditory nerve: *t*_(12)_ = 4.3, CF of the hippocampi: *t*_(12)_ = 4.0, CF of the amygdalae: *t*_(12)_ = 4.1) and from TP1 to TP3 (right TR of the auditory nerve: *t*_(12)_ = 3.9, CF of the hippocampi: *t*_(12)_ = 4.6, CF of the amygdalae: *t*_(12)_ = 4.4). In AD and MD, both indices were significantly different (AD: *F*_(2, 24)_ = 6.1; *p* = 0.0073, MD: *F*_(2, 24)_ = 5.9; *p* = 0.0081) in the segments of the right UF over three time points. The post hoc analysis revealed that the segments significantly decreased from TP1 to TP2 in both AD and MD (AD: *t*_(12)_ = 4.6, MD: *t*_(12)_ = 4.4). From TP1 to TP3, only AD significantly decreased (AD: *t*_(12)_ = 2.2), whereas MD marginally decreased (MD: *t*_(12)_ = 2.1, *p* = 0.0615). These differences are presented as percent changes in Figure 7.

**Fig. 7.**
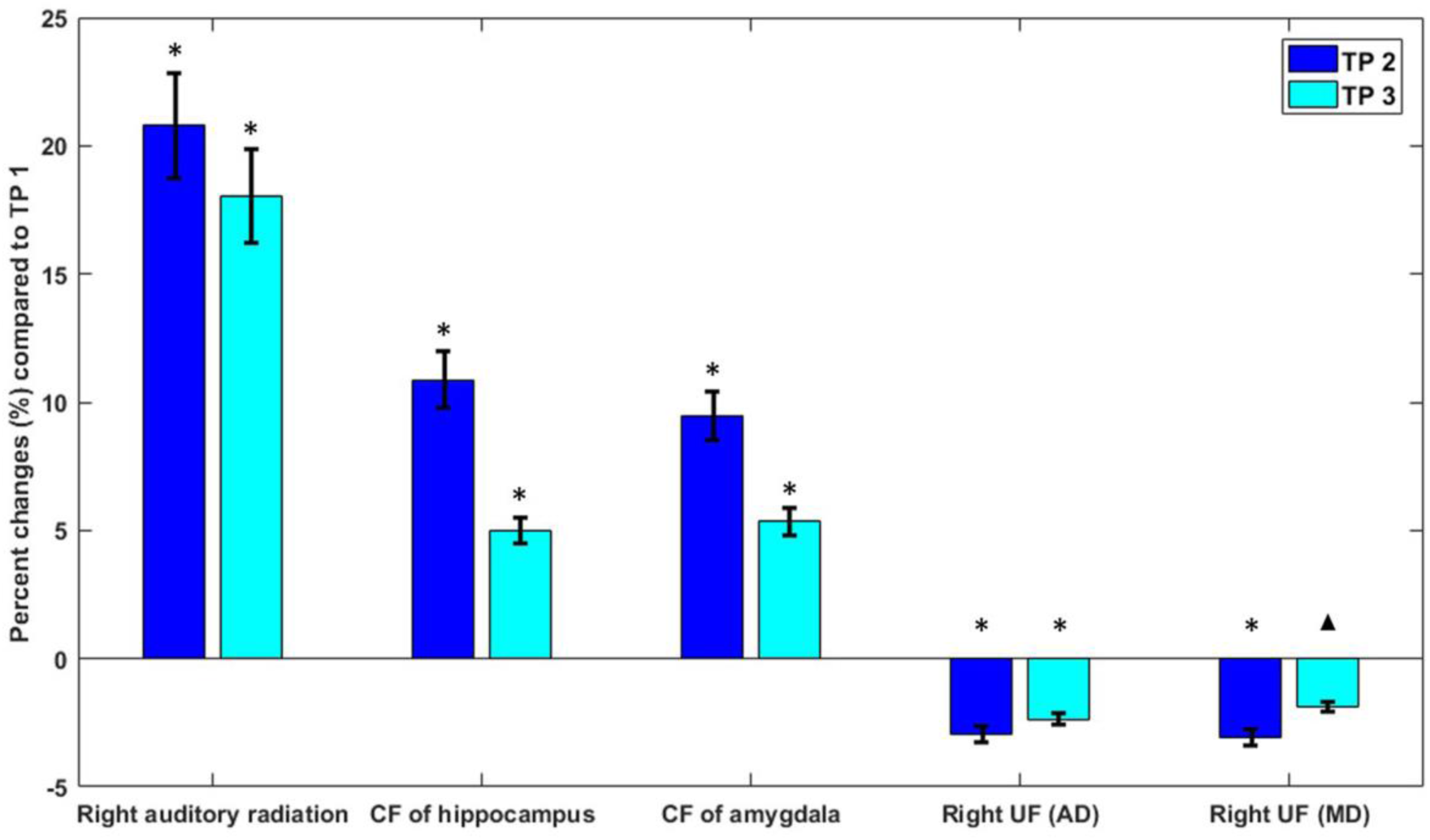
Percent changes in the tract segments that indicate sustained modulation after MBSR training. The percent changes between TP2 and TP1 (baseline) and between TP3 and TP1 are displayed as dark blue and light blue bars, respectively. The GFA values in the right TR of the auditory nerve, CF of the hippocampi, and CF of the amygdalae reveal significant diffenerces compared with the baseline. In the right UF, AD exhibits a sustained difference until TP3, whereas MD exhibits a marginal trend of difference. The error bar indicates one standard error of percent change. The asterisk and triangle indicate the significant and marginal differences, respectively.

**Table 2.** Results of repeated measures ANOVA in the 6-month follow-up study

| Diffusion indices | Tract | MBSR group |  |  | Main effect |  | Pair-wise |  |
| --- | --- | --- | --- | --- | --- | --- | --- | --- |
|  |  | Time point 1<br>Mean (SD) | Time point 2<br>Mean (SD) | Time point 3<br>Mean (SD) | *F* | *p* value | 1 vs 2 | 1 vs 3 |
| **GFA** | **R TR of auditory nerve** | 0.273 (0.040) | 0.328 (0.054) | 0.319 (0.046) | 11.1 | < 0.001 | < 0.001 | < 0.01 |
|  | **CF of hippocampus** | 0.547 (0.073) | 0.606 (0.090) | 0.575 (0.081) | 13.0 | < 0.01<sup>#</sup> | < 0.01 | < 0.001 |
|  | **CF of amygdala** | 0.381 (0.052) | 0.417 (0.063) | 0.401 (0.051) | 13.1 | < 0.001<sup>#</sup> | < 0.01 | < 0.001 |
| **AD** | **R UF<sup>+</sup>** | 7.729 (0.257) | 7.498 (0.237) | 7.543 (0.316) | 6.1 | < 0.01 | < 0.001 | < 0.05 |
| **MD** | **R UF<sup>+</sup>** | 5.327 (0.291) | 5.162 (0.295) | 5.221 (0.246) | 5.9 | < 0.01 | < 0.001 | 0.0615 |

<sup>+</sup>: ( $\times 10^{-4}$ ) mm<sup>2</sup>/sec unit
<sup>#</sup>: Greenhouse-Geisser correction for violation of sphericity
Note: Time point 1: pre-MBSR scan
Time point 2: post-MBSR scan
Time point 3: follow-up scan

## Discussion

Growing evidence suggests that the brain can be modulated through meditation practices. Most studies in this field have adopted a “Pre–Post” longitudinal design to examine the difference in brain structures before and after short-term meditation training. However, whether the changes induced by short-term meditation training are transient or sustained after a period of cessation remains unclear. In the present study, we longitudinally examined the effects of an 8-week MBSR training program on the microstructural properties of the white matter in meditation-naïve participants. The study entailed three scans at three time points; the first scan was performed before the MBSR program, the second immediately after the completion of the program, and the third 6 months after the completion of the program (with cessation of meditation practice during this period). We observed an increase in GFA values from TP1 to TP2 in the right TR of the auditory nerve, CF connecting the bilateral amygdalae, and CF connecting the bilateral hippocampi, and a sustained increase in GFA values at TP3 compared with TP1. In addition, a decrease in AD and MD was observed at TP2 and TP3 in the right UF. Our results reveal that the 8-week MBSR training program induced modulations in the microstructural properties of the white matter that were sustained even after practice was discontinued.

MBSR training is a mental skill acquisition technique aimed at instructing trainees to be nonjudgmental and aware of present-moment experience. Modern mental training programs such as MBSR, mindfulness-based cognitive therapy, and IBMT are packaged into a short-term training course to assist practitioners in learning the core values of mindfulness meditation and in effectively and systematically practicing these skills. Short-term MBSR training emphasizes the need for mental effort and conscious control to achieve a meditative state (Tang et al. 2012a). As such, practitioners learn to focus on a target of awareness, such as breathing, body sensation, or other present-moment experience perceived during open monitoring. The core values of the 8-week MBSR program include the self-regulation of attention, and, more importantly, a nonjudgmental attitude toward present-moment experience (Bishop 2004). Accordingly, this mindfulness practice may affect practitioners’ psychological state by altering their perspectives on internal experience. Individuals who undertake this mindfulness practice learn to observe their sensations, thoughts, and emotions in an objective manner and focus on the process rather than the content of this awareness. By doing so, individuals can develop insight into their customary proclivities of thinking, allowing them to change negative patterns of thinking and react differently from their habitual manner (Robins et al. 2012). This learning process is believed to allow individuals to engage in emotional learning and regulation, which is the main psychological change derived from MBSR (Goldin and Gross 2010; Roemer et al. 2015). Following the 8-week MBSR training program, practitioners can improve their cognitive, perceptual, emotional, or even motor performance through mindfulness skill learning. A previous randomized controlled trial on patients with anxiety disorders demonstrated that the mindfulness-related cognitive effects persisted 6 months after an 8-week MBSR program (Vollestad et al. 2011). These findings indicate that mindfulness training has sustained beneficial effects on disease-related symptoms compared with the control condition.

Mental learning and training have been demonstrated to modulate the gray matter and white matter (Chapman et al. 2015; Saleem and Samudrala 2017; Zatorre et al. 2012). Tang et al. (2010) reported that 11 h of IBMT can induce changes in the FA values of the white matter tracts connecting the anterior cingulate cortex in meditation novices. In another study conducted by Mackey et al. (2012), the participants undertook 3 months of preparation for the Law School Admission Test, which requires strong reasoning skills. Altered radial and mean diffusivities were observed in the white matter tracts connecting the frontal and parietal cortices. These findings suggest that short-term intensive skill training or learning modulates the microstructural properties of the relevant white matter tracts. The current study showed that such microstructural changes can be sustained without continual practice for at least 6 months after completion of an 8-week MBSR training program. According to our review of the relevant literature, this is one of the first studies to address the sustainability of microstructural changes induced by mental training. In other activities, it was reported that the structural neuroplasticity induced by short-term motor training was sustained without continual practice. Scholz et al. designed a 6-week training program to train novices to juggle (Scholz et al. 2009). The participants were scanned using DTI to measure the white matter change before and after the training program and after a 4-week follow-up period without practice. They observed a significant training-induced increase in the FA values in the white matter underneath the right posterior intraparietal sulcus. Notably, the change in the training group remained elevated after the 4-week follow-up period. Draganski et al. (2006) investigated gray matter change following a period of intensive learning for a medical examination and found that the volume of the posterior hippocampus remained significantly increased even after a semester break of 3 months. These studies indicate that neuroplasticity might be invoked to meet the need for processing novel informational demands (Chambers et al. 2004), and the subsequent effect on brain structures is still detectable even when the participants no longer continue to practice.

The brain is a dynamic system in which neuroplasticity plays an important role. Neuroplasticity can be induced by motor and sensory processing and by more complicated mental activities such as meditation training (Fox et al. 2014). The 8-week MBSR training applied in this study is a systematic mental training program that involves process-specific learning, a type of learning that can be generalized to novel stimuli or task contexts (Slagter et al. 2011). Although the exact neurological mechanism underlying the lasting effect of MBSR on structural change is still unclear, we propose that the sustained alteration of the brain structures might be related to process-specific learning such as emotional learning.

In the current study, we found that the microstructural properties of the white matter were significantly modulated after the 8-week MBSR training program. Previous studies have reported white matter changes induced by short-term mindfulness meditation training in adults (Kang et al. 2013; Posner et al. 2014; Tang et al. 2012b, 2010). Tang et al. (2012b) found increased FA values in the corona radiata following a 4-week IBMT program and also reported reduced RD and AD in the corpus callosum, corona radiata, and superior longitudinal fasciculus. Increased myelination was associated with increases in the FA values after motor skill learning in an animal study (Sampaio-Baptista et al. 2013). Furthermore, increased FA values were correlated with an increased ratio of the myelinated axons (Yamasaki et al. 2015). Therefore, in the present study, we speculate that the increased GFA values in the CF connecting the bilateral hippocampi and amygdalae and the right TR of the auditory nerve might be related to the increased myelination induced by short-term MBSR training. Moreover, a reduced AD may reflect an increase in axonal density or caliber (Suzuki et al. 2003) and might be one of the mechanisms underlying the decrease in the AD value in the right UF in the present study. In sum, after the 8-week MBSR training program, the white matter connection was improved in the right TR of the auditory nerve, CF connecting the bilateral hippocampi, CF connecting the bilateral amygdalae, and right UF.

Previous studies have reported structural differences in the hippocampal formation in meditators (Fox et al. 2014; Luders et al. 2013). The hippocampus appears to be critical for contextualizing emotional learning (i.e., facilitating emotional responses that consider the current context; Morgan et al. 1993). The results of the present study indicate that in addition to morphological changes in the hippocampus, the connectivity of the bilateral hippocampi is improved through the CF. Improved connectivity of the bilateral amygdalae was also observed in our study. Holzel et al. (2010) reported that MBSR-induced stress reduction was associated with a reduced volume of the right amygdala. The amygdala is an important limbic structure related to emotional state. It receives sensory information and projects it to other subcortical structures, mediating emotion-related physiological and behavioral effects. Structural differences in the corpus callosum have been reported in both long-term and short-term meditation practitioners (Luders et al. 2012; Tang et al. 2010). These results and ours suggest that alterations of the hippocampal and amygdala CF might be the consequences of an enhanced connection between the bilateral hemispheres in memory and emotion.

In addition to the increased connections of the bilateral hippocampi and amygdalae, an enhanced connection between the right primary auditory cortex and thalamus was indicated by an increased GFA value after MBSR training. Previous studies have demonstrated that MBSR practitioners exhibit altered functional connectivity between the anterior default-mode network and auditory network, which indicates greater reflective awareness for auditory experiences (Kilpatrick et al. 2011). Recent studies have further suggested that the auditory cortex not only processes auditory information but is also responsible for emotional learning (Grosso et al. 2015). In addition to processing and encoding the physical attributes of the conditioned tones, the auditory cortex, including the primary and higher orders, changes auditory acuity in response to emotional experiences. The auditory cortex was found to be involved in all the phases of emotional memory processing, from learning to the remote storage and retrieval of information (Grosso et al. 2015). Therefore, we hypothesize that improved connection in the TR of the auditory nerve is associated with emotional regulation, not merely the awareness of external acoustic stimuli.

Changes in the UF have been observed following meditation training (Kang et al. 2013; Luders et al. 2011). The UF is involved in emotional learning and regulation and is considered to transmit salience-laden stimuli stored in the anterior medial regions of the temporal lobe to the lateral orbital frontal cortex (Von Der Heide et al. 2013). Holzel et al. (2016) reported that the right UF exhibited a significant increase in the FA value following short-term MBSR training. Another study reported that patients with right hemisphere stroke involving the right UF exhibited significantly impaired function in an empathy task compared with patients without lesions in the right UF (Oishi et al. 2015), suggesting that the right UF may be involved in the empathy network. The present study found a significant decrease in AD in the right UF following MBSR, which might reflect an increase in axonal density or caliber (Suzuki et al. 2003). Although our diffusion index with a significant difference is not the same as that reported previously, it is consistent with the notion that the connectivity in the right UF is increased following MBSR training. In sum, our findings suggest that the changes in white matter tract properties following an 8-week MBSR training program are associated with emotional regulation and emotional learning. Because the neural substrates plasticized by short-term MBSR training are involved in emotion-related networks, these changes can facilitate the processing of emotional experiences.

This study has some limitations. First, the control group mitigated the effects of some confounding variables, such as age, sex, and other demographic factors; these participants did not receive any psychological intervention. An alternative active control design is suggested to provide control participants with active interventions such as relaxation training or stress management education. This active control design can control for confounding factors such as amount of physical exercise, interaction with other participants, and psychoeducation, and can extract meditation-specific effects to assess the unique underlying mechanisms of mindfulness meditation practice. Second, the sample size of the MBSR group was relatively small compared with that in other studies, which might have reduced the statistical power of the present study. Our study was longitudinal and involved three scans at three visits over a span of 8 months. Under such circumstances, completion of the whole study by all participants is very difficult to attain, as evidenced by our low success rate (43%); 17 out of 30 participants dropped out. Third, we failed to identify any significant correlations between psychological improvements and the observed changes in white matter integrity. Given that mindfulness meditation influences a wide spectrum of psychological performance, we may have failed to assess the core values of mindfulness meditation, such as emotion, memory, and sensation, which are relevant to the observed alteration in white matter integrity. Another possible reason for the negative finding is that most of the structural changes were related to the amount of time spent on training rather than the training outcome (Scholz et al. 2009). Future research should investigate the relationship between structural changes and emotional learning induced by mindfulness practice. A larger sample size and an active control group may be an ideal paradigm for investigating training-induced neuroanatomical effects. If confirmed by future research, the findings described here might be highly relevant to the neurological mechanisms of mindfulness meditation and clarify the process through which mindfulness exerts its beneficial effects on mental health and well-being.

The findings of the present study suggest that sustained modulation of the altered white matter integrity is observed even without the continuation of the mindfulness meditation practice for 6 months after the 8-week MBSR program. The enhanced white matter tracts (i.e., the right auditory radiation, CF connecting the bilateral amygdalae, CF connecting the bilateral hippocampi, and right UF) might be the neural substrates of improved emotional learning and regulation through the modulation of connections between gray matter regions involved in the emotion network. Although the underlying neurological mechanism has not yet been validated, neuroimaging studies suggest that strengthening of white matter integrity might be related to experience-dependent neuroplasticity. According to the review of previous studies and our results, the microstructural changes in the white matter tracts following MBSR training may improve emotional learning and regulation. Sustained neuroplasticity in the microstructural properties of the white matter provides physical evidence of a prolonged learning effect induced by mental training. Further research is warranted to obtain unequivocal results elucidating the time course of training-related microstructural changes in the white matter.

## Acknowledgments

We cordially thank all participants for their contribution and partaking in the study. We also thank Ruentex Group for their funding support. This manuscript was edited by Wallace Academic Editing.

## Research involving Human Participants and/or Animals

This research only involved healthy human participants.

## Ethical approval

All procedures performed in this study involving human participants were in accordance with the ethical standards of the National Taiwan University Hospital (NTUH) Research Ethics Committee (REC) and with the 1964 Helsinki declaration and its later amendments or comparable ethical standards. (NTUH-REC approve number: 201412013RIFC)

## Informed consent

Informed consent was obtained from all individual participants included in the study.

